# Spatial Metabolome Lipidome and Glycome from a Single brain Section

**DOI:** 10.1101/2023.07.22.550155

**Authors:** Harrison A. Clarke, Xin Ma, Cameron J. Shedlock, Terrymar Medina, Tara R. Hawkinson, Lei Wu, Roberto A. Ribas, Shannon Keohane, Sakthivel Ravi, Jennifer Bizon, Sara Burke, Jose Francisco Abisambra, Matthew Merritt, Boone Prentice, Craig W. Vander Kooi, Matthew S. Gentry, Li Chen, Ramon C. Sun

## Abstract

Metabolites, lipids, and glycans are fundamental biomolecules involved in complex biological systems. They are metabolically channeled through a myriad of pathways and molecular processes that define the physiology and pathology of an organism. Here, we present a blueprint for the simultaneous analysis of spatial metabolome, lipidome, and glycome from a single tissue section using mass spectrometry imaging. Complimenting an original experimental protocol, our workflow includes a computational framework called Spatial Augmented Multiomics Interface (Sami) that offers multiomics integration, high dimensionality clustering, spatial anatomical mapping with matched multiomics features, and metabolic pathway enrichment to providing unprecedented insights into the spatial distribution and interaction of these biomolecules in mammalian tissue biology.

## INTRODUCTION

Metabolomics (Fiehn, 2002; Gibney et al., 2005; Lisec et al., 2006), lipidomics (Cajka and Fiehn, 2016; Han and Gross, 2005), and glycomics (Cummings and Pierce, 2014; Ruhaak et al., 2010; Wada et al., 2007) are three distinct facets of omics methodologies, each offering a unique window into the connected and complex biochemical processes in living organisms. The current state of these fields lacks spatial resolution and unified, integrated analyses that offer a broad overview of the interconnected metabolic landscape. The development of an integrated spatially-resolved metabolomics, lipidomics, and glycomics is crucial for advancing our knowledge of biological systems and has the potential to transform our understanding of the complex tissue metabolic heterogeneity, uncover novel biomarkers and even therapeutic targets. Nevertheless, the development of such integrated approaches is challenged by the inherent differences in physicochemical properties and analytical requirements of each molecular class.

Matrix-assisted laser desorption/ionization (MALDI) mass spectrometry imaging emerged as a powerful tool for spatially-resolved molecular analysis, offering the possibility to overcome major limitations associated with pooled sample analysis (Caprioli et al., 1997; McDonnell and Heeren, 2007). Individual developments in spatial-metabolomics (Wang et al., 2015), -lipidomics (Djambazova et al., 2020; Zemski Berry et al., 2011), and -glycomics (Conroy et al., 2023; Drake et al., 2017; Powers et al., 2014; Velickovic et al., 2022; Wei and Li, 2009) methodology are underway even at the single cell level (Rappez et al., 2021). To this end, recent advances in MALDI imaging have enabled multiplexed analysis of diverse biomolecules (Black et al., 2019), such as the co-analysis of N-linked glycans and extracellular matrix proteins, or N-linked glycans and storage carbohydrates such as glycogen by optimizing sample preparation, enzyme application, matrix selection, and instrumental parameters (Clift et al., 2018; Clift et al., 2021; Young et al., 2022). These developments have poised MALDI imaging for broad adaptation in metabolism research, as they facilitate the acquisition of comprehensive and spatially-resolved molecular data. Further, the complexity and high dimensional nature of the datasets calls for the need of a robust computational pipeline to extract actionable and biologically relevant information for hypothesis generation (Conroy et al., 2023). An integrated workflow combining metabolomics, lipidomics, and glycomics from a single tissue section would offer unprecedented insights in to the spatial and heterogenous metabolic landscape of mammalian tissues, driving the next wave of metabolism research in health and diseases.

In this study, we present a comprehensive roadmap for the integration of spatial metabolomics, lipidomics, and glycomics from a single tissue section. We demonstrate a sequential sample preparation strategy circumvents cell-loss through adjacent tissue sections as well as batch effects in different omics domains. This method enables the acquisition of high-resolution, spatially-resolved data for broad biomolecule classes. Furthermore, we implemented a robust computational framework to integrate these datasets, enabling the identification of spatial patterns and functional relationships between metabolites, lipids, and glycans. Our approach demonstrates the significant impact of integrated spatial omics on our understanding of complex biological systems, revealing previously unrecognized molecular features and associations within mammalian tissues.

## RESULTS

### A workflow for spatial triple-omics by mass spectrometry imaging

The application of spatial metabolome, lipidome, and glycome on a single tissue section would presents a multitude of benefits, primarily in maintaining the spatial continuity of the original tissue microenvironment and conserving tissue resources (Bartolacci et al., 2022; Cornett et al., 2007; Duhamel et al., 2022; Sun et al., 2023). A tissue multi-omic approach facilitating a more holistic understanding of metabolic underpinnings of tissue metabolism and biology. Using a mouse brain as a model, we successfully acquired multiomics data from a single tissue section (**Figure. 1**). A single 10 µm thick brain section coated with N-(1-naphthyl) ethylenediamine dihydrochloride (NEDC) matrix was first subjected to sequential spatial metabolome and lipidome scans using MALDI mass spectrometry imaging (Figure. 1A). NEDC matrix and negative ionization mode are amenable for both spatial metabolome (Chen et al., 2012) and lipidome (Wang et al., 2018) analyses. Following spatial lipidome imaging, the NEDC matrix was removed, and the tissue was fixed, then by Peptide - N -Glycosidase F (PnGase F) and Isoamylase digestion to release complex carbohydrates for spatial glycome and glycogen analysis (Hawkinson et al., 2022; Hawkinson and Sun, 2022). Spatial glycomics was performed in positive mode using α-cyano-4-hydroxycinnamic acid (CHCA) as the ionization matrix (**Figure. 1A**). We successfully demonstrated acquisition of high-quality spatial images for molecular features in all three omics classes (**Figure. 1B** and **Supplemental Figure 1**). These include spatially unique metabolites, lipids, and glycans among different regions of a coronally cut mouse brain (**Figure. 1B** and **Supplemental Figure 1**). This method allows for a more holistic perspective of the metabolic landscape of the tissue and seamlessly combines three omic analyses on a single tissue section, thereby preserving the spatial congruity.

**Figure 1.**
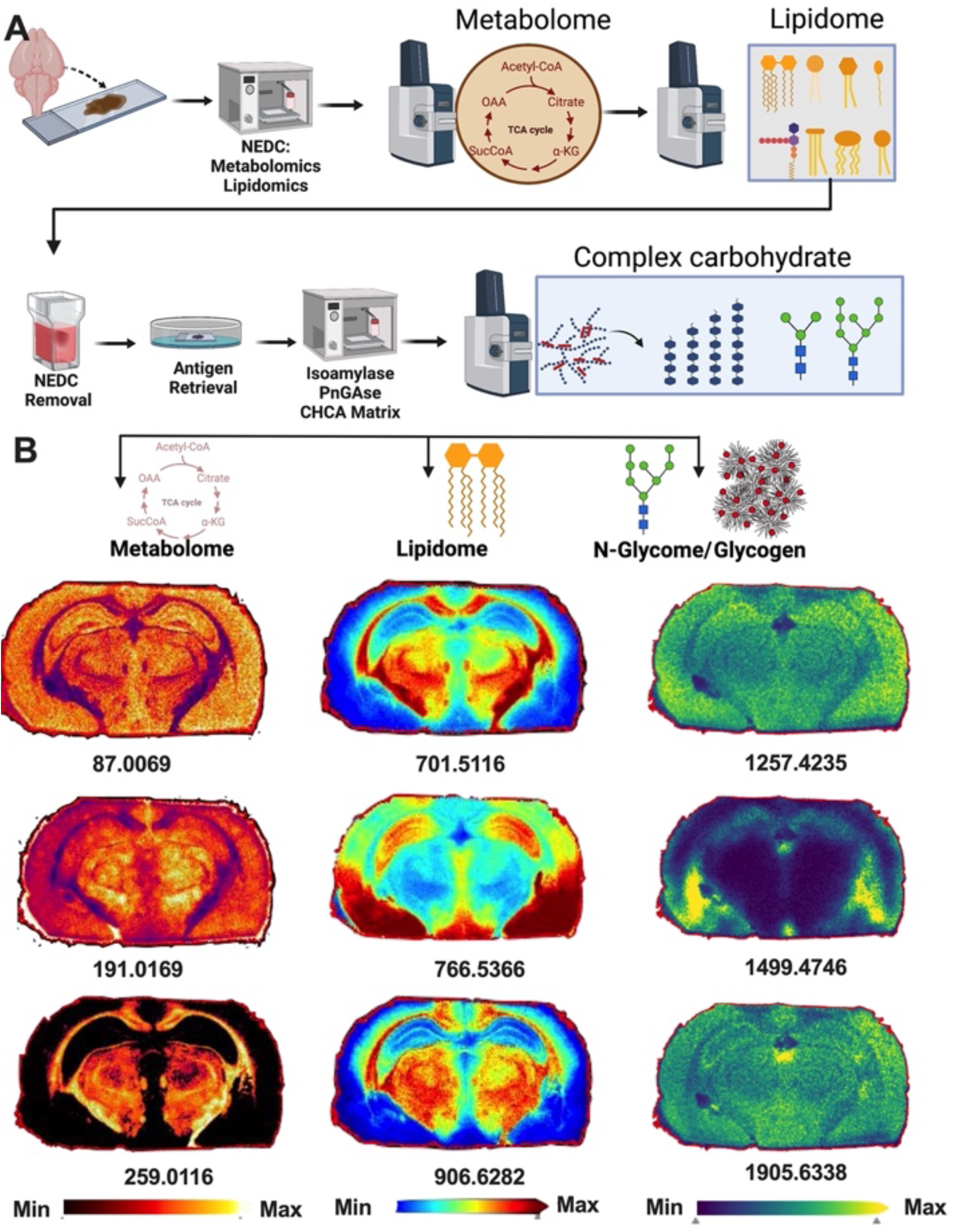
workflow for simultaneous spatial metabolome, lipidome, and glycome by mass spectrometry imaging. **a** Schematics workflow for the simultaneous spatial assessment of metabolome, lipidome, and glycome by sequential MALDI imaging. Fresh frozen mouse brain were sectioned to 10um thickness followed by application of NEDC matrix for the MALDI imaging of metabolites and lipids. The same tissue section is then prepared for isoamylase and PnGase F treatment followed by CHCA matrix application and MALDI imaging of complex carbohydrates such as N-glycans and glycogen. Created with BioRender.com. **b** Spatial heatmap images of selected biomolecules from each of metabolome, lipidome, and glycome MALDI imaging runs. All images are produced from the same tissue section following the protocol outlined in a. *m/z* values are below top of the heatmap.

### Co-registering and integration of Spatial multiomics datasets

With >10^5^ pixels scanned per biological sample, the translation of MALDI multiomics datasets into actionable information to aid hypothesis-driven research presents a significant computational challenge. Recognizing this critical need, we developed a bioinformatic pipeline termed Spatial Augmented Multiomics Interface (Sami) that performs multiomics integration, high dimensionality reduction and clustering, spatial clustering, annotation, and pathway enrichment in one platform for the comprehensive analysis of spatial metabolome, lipidome, and glycome datasets (**Figure. 2A**). The first step of Sami is to perform multiomics integration, which consolidates the disparate biological features of metabolomics, lipidomics, and glycomics data into a singular, harmonized input suitable for high-dimensionality and pathway analyses (**Figure. 2A**). The smartbeam laser technology within the MALDI instrument registers precision coordinates of the laser shots and preserve a unified x and y coordinates of each pixel across each omics modalities. By cross-referencing the *x* and *y* coordinates, Sami processing can determine the exact spatial location of each pixel in all datasets, effectively superimposing these multiple layers of omics information on a single spatial metadata location. Successful integration opens the opportunity for additional analyses such as correlation and network analyses across modalities. We found a number of metabolome and lipid features shown high levels co-expression, their abundance and spatial distribution (**Figure. 2B-C** and **Supplemental Figure** 2). For example, 256.995 *m/z* from metabolomics and 862.606 *m/z* from the lipidomics analysis exhibit high levels of spatial co-expression identified after multiomics integration (**Figure. 2B)**. Finally, we demonstrated multiomics integration through inter and intra domain connectivity via network analysis (**Figure. 2D**). Network analysis demonstrate robust connectivity between molecule classes, future support that notion that biomolecules with different physiochemical properties are metabolically channeled. (**Figure. 2D**). Collectively, our data support the generation of a robust, integrated, single dataset from three different omics modalities.

**Figure 2.**
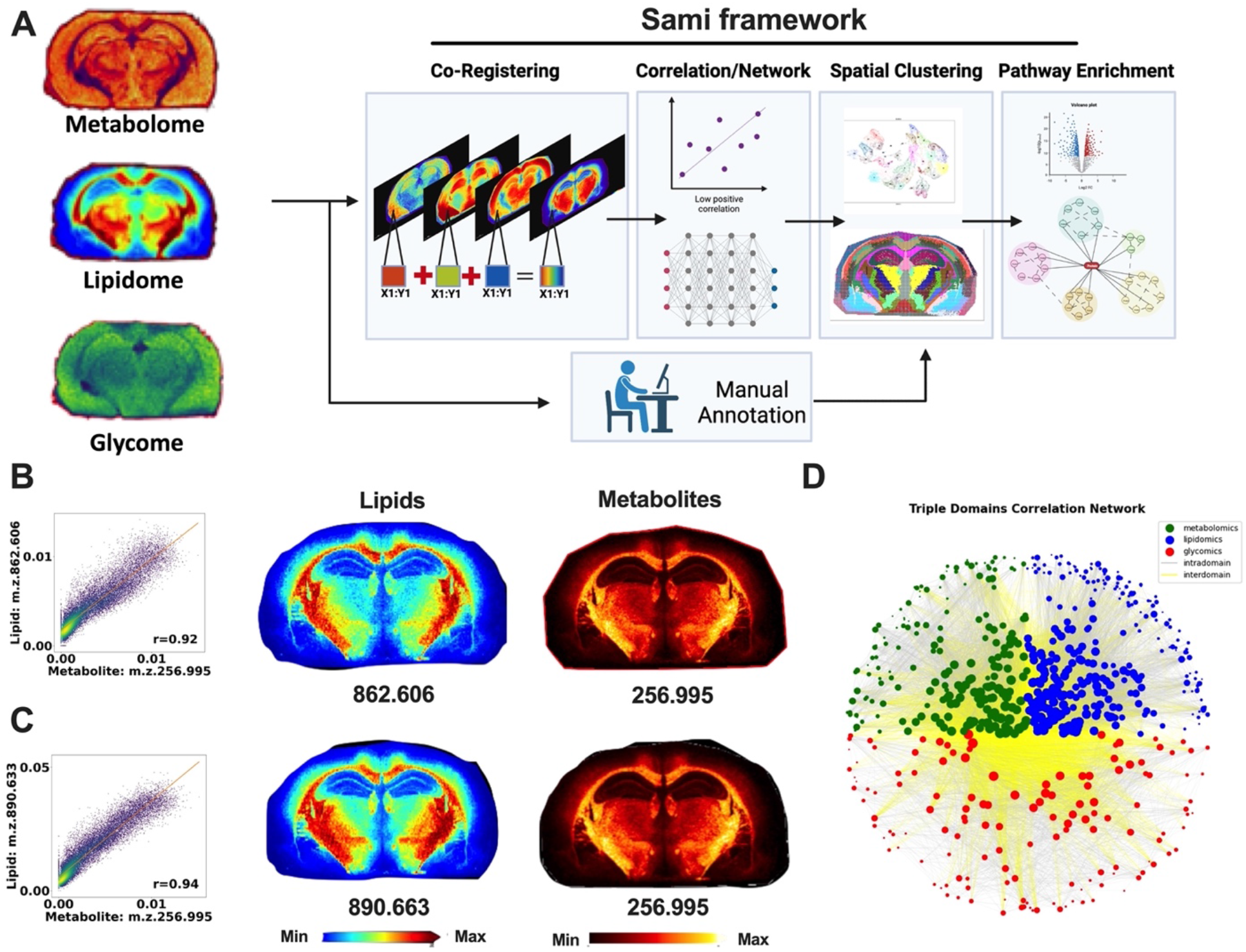
Spatial Augmented Multiomics Interface (Sami). **A**. Computational framework for Spatial Augmented Multiomics Interface (Sami). B and C. Scatterplots showing correlation between metabolome and lipidome datasets based on Pearson correlation coefficient after multiomic integration. Paired spatial heatmap images are shown on the right. All images are produced from the same tissue section, *m/z* values are below the heatmap. **d** Network analysis of the integrated multiomics dataset demonstrating intra and interdomain connectivity. Intradomain is within each omic dataset, and interdomain connectivity represents correlations among different omics modalities. Metabolome (green), lipidome (blue), glycome (red) after multiomics integration. Intra domain connections are presented as grey lines and interdomain are presented as yellow lines.

### High dimensionality reduction, spatial clustering, and annotation

High-dimensionality reduction as part of the Sami framework serves as a powerful strategy to distill high dimensional MALDI MSI datasets to actionable and manageable information (**Figure 3A**). Upon performing the spatial clustering, an intriguing observation was made – the spatial clusters exhibited a striking correspondence with the anatomically distinct brain regions (Dong, 2008; Sunkin et al., 2012) (**Figure 3B**). This is particularly transformative as it demonstrates the capacity to leverage and translate spatial information within the multiomics data to biology. To further validate this finding, we performed manual annotated of each spatial cluster against the brain regions represented by the Mouse Brain Atlas from the Allen Institute (Sunkin et al., 2012) (**Figure. 3B-C**). This rigorous comparison revealed a high degree of similarity between the spatial clusters and the recognized brain regions, offering compelling evidence of the accuracy and biological relevance of UMAP clustering that relates to the biochemistry of the brain. To further enhance the utility of the spatial multiomics map, we then established a reference multiomics feature set for the annotation of brain regions. Selected features for represented clusters are shown as Radial trees and density plots (**Supplemental Figure. 3A-B**). Annotated clusters can serve as a reference guide, aiding in the assignment and interpretation of multiomics features across different studies. To confirm the robustness of the annotations derived through Sami, we performed spatial triple-omics in a separate brain section (test brain) and performed unsupervised spatial clustering. Following, each cluster in the test brain were matched to the original reference brain section through feature matching to create a supervised high dimensional cluster matching the reference brain, therefore can be annotated for brain regions (**Figure. 3D-E**). The results obtained corroborated our initial findings, with an excellent match between the reference and test brain datasets. This feature will serve as foundation for future brain-related researches using spatial multi-omics.

**Figure 3.**
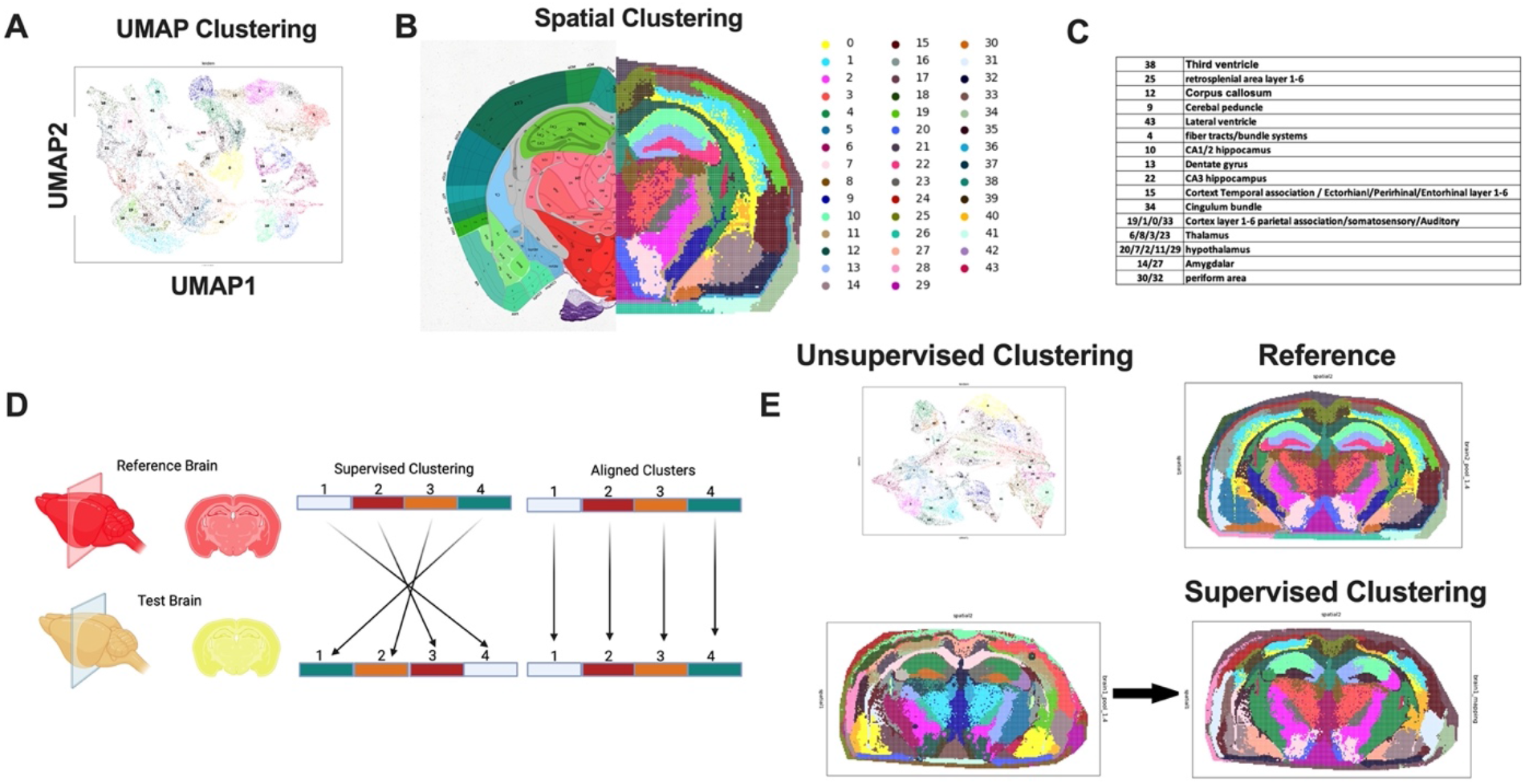
Application of high dimensionality reduction, spatial clustering, and manual annotation. **A**. High dimensionality reduction and UMAP plot of all available clusters. **B**. Spatial mapping of each clusters based on x and y coordinates of MALDI laser spots. Right half of the spatial clustering map is merged with the left side of brain regions adapted from Allen Mouse Brain Atlas, (mouse.brain-map.org and atlas.brain-map.org), color and their corresponding cluster numbers are on the right side. **C**. cluster number and their annotation based on matching with the Allen Brain Atlas. D. Schematics of supervised cluster match in a new brain section. **E**. Supervised spatial clustering from a reference brain. A second brain undergone spatial metabolome, lipidome, and glycome MALDI imaging following high dimensionality reduction and supervised spatial clustering by annotation and matching clusters with reference brain based on over represented features.

### Biological insights through pathway enrichment

Subsequent to spatial clustering and annotation, Sami performs pathway analysis, a key step to provide biological insights into the metabolic heterogeneity of distinct clusters or anatomical brain regions. First, we performed differential expression analysis by comparing each cluster with the mean of the rest of the clusters as illustrated by volcano plots (**Figure. 4A & Supplemental Figure. 4**); From differential expression analysis, the Top 50 annotated multiomics features from own library (Conroy et al., 2023; Conroy et al., 2021b) and previously published assignments (Wang et al., 2022a; Wang et al., 2015; Wang et al., 2022b) (**Supplemental Table. 1**) of each cluster exhibiting significant alterations were selected for metabolic pathway enrichment analysis via the MetaboAnalyst 3.2 pipeline embedded in Sami. We conducted separate enrichment analyses: one for metabolomics and glycomics via the Small Molecule Pathway Database (SMPDB) (Frolkis et al., 2010), and another for lipids using the lipid library integrated within MetaboAnalyst (**Supplemental Figure. 5**). As an example, metabolic pathways enriched for the CA3 region of the mouse brain are “mitochondrial electron transport chain”, “transfer of acetyl groups to mitochondria”, “glycolysis”, and “purine metabolism” (**Figure. 4B**). In contrast, the cortex showed unique enrichment for “aspartate metabolism” and “sphingolipid metabolism” (**Supplemental Figure. 5**). By correlating these pathways with specific brain regions and clusters, we underscore the utility of Sami in mapping the regional metabolic diversity within the brain. Collectively, Sami paves the way for a spatially resolved understanding of the complex multiomics landscape of the brain, and potentially other complex biological systems.

**Figure 4.**
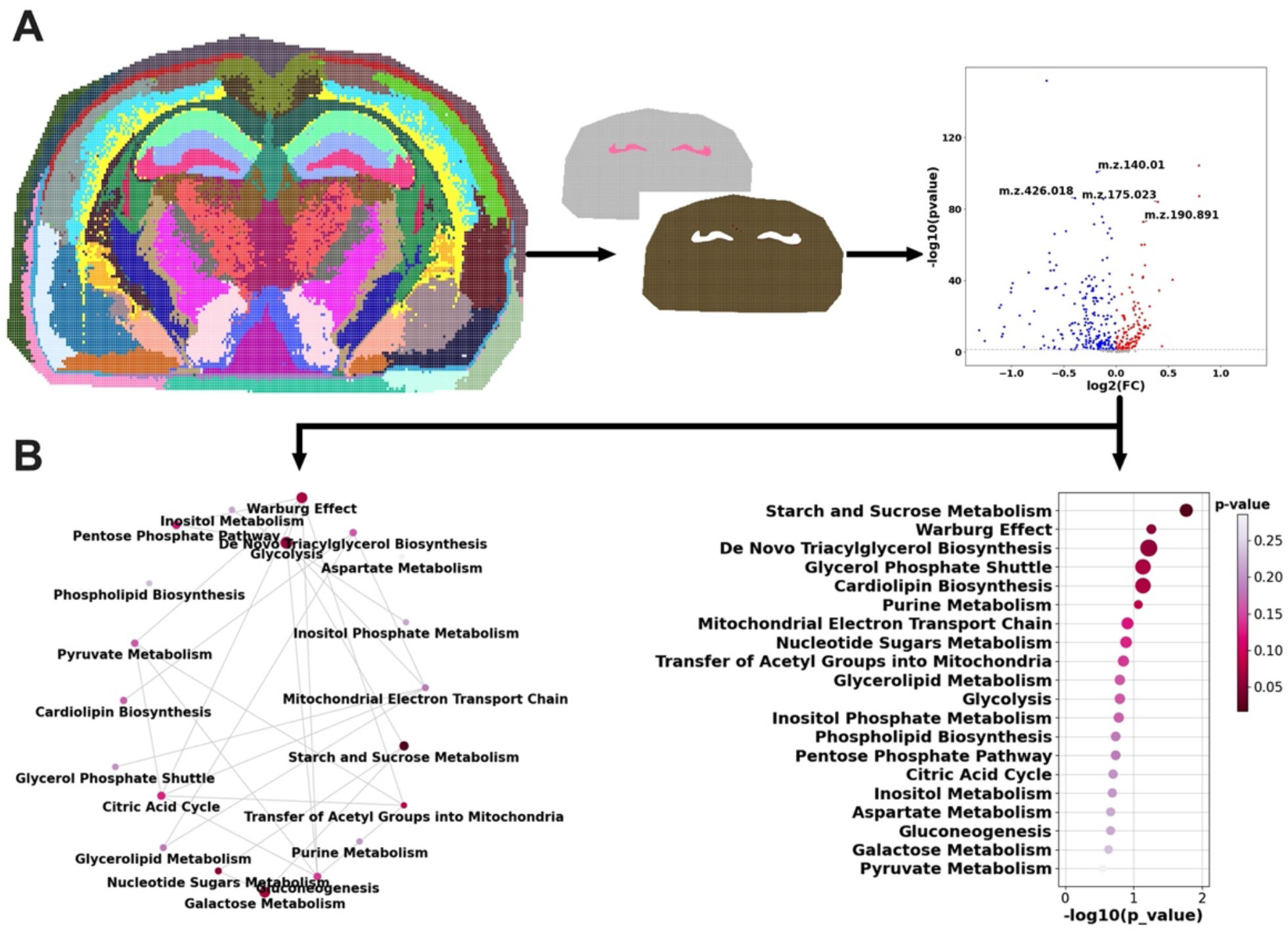
Pathway enrichment within the Sami pipeline. **A**. Schematics showing pathway enrichment of of the CA3 region of the mouse brain. Volcano plot for metabolome dataset is shown significant changes between CA3 and rest of the brain regions. P value greater than 0.05 are considered significant. Increased features are in red and decreased features are in blue. **B**. Metabolic network plot and metabolic pathway enrichment based on enriched metabolic pathways generated from the Metaboanlyst 3 R package embeded in Sami.

## Discussion

Metabolomics, lipidomics, and glycomics are corner stone techniques driving biological discovery in the last several decades (Hart and Copeland, 2010; Wenk, 2005; Worley and Powers, 2013). In this study, we present a roadmap to perform simultaneous assessments of the metabolome, lipidome, and glycome from a single tissue section. This represents a major advancement compared to traditional pooled omics analyses, providing a broad and comprehensive assessment of the metabolic landscape. Our approach overcomes several limitations of conventional methods, offering insights into the spatial distributions of a wide array of biomolecules, thereby fostering a nuanced understanding of the metabolic processes within complex tissue microenvironments. In addition, Sami presents an improved level of rigor and reproducibility by minimizing sample handling and fractionation steps (Smith et al., 2020). The entire pipeline can be completed within a span of 48 hours when imaging at a resolution of 50µm, Sami offers a promising platform for high-throughput omics studies for the future.

In addition, we also report the robust computational framework of Sami, a series of tools designed specifically to handle the vast amount of data produced by multiomics MALDI imaging runs. Sami’s ability to integrate and reduce high-dimensional data into actionable datasets represents a leap forward in the quest to understand biological systems and foster future hypothesis-driven research. One salient finding from our study is the efficacy of metabolism as a classifier of brain regions, a concept that holds immense implications for our understanding of brain functionality and neurological disorders (Camandola and Mattson, 2017; Conroy et al., 2021a; Sun et al., 2021). The differential metabolic profiles unveiled through our pathway enrichment analyses, demarcate specific brain regions, emphasizing the spatial heterogeneity of metabolic processes within the brain. These findings hold the potential to revolutionize our understanding of brain metabolism, paving the way for novel research avenues into the complex interplay between brain function and metabolic activity.

The current study represents the inaugural version of our computational workflow. Future development will focus on expanding our molecular library (Wadie et al., 2023) and improving laser raster size of 5µm, which is near single-cell level (Guenther et al., 2011; Yang et al., 2014). We anticipate 5µm multiomics imaging would offer deeper insights into metabolic cross talk and heterogeneity at the single-cell level. Additionally, while our current annotations focus on the brain, future work should seek to identify metabolic classifiers for other tissues, such as liver, kidney, and lung, and extend these analyses to metabolic diseases such as cancers, neurological disorders, and inborn errors of metabolism. The Sami framework is designed to accommodate these updates, functioning as an “expandable” system that can be seamlessly updated as new improvements emerge. In conclusion, our data show that Sami is a powerful tool for multiomics research; we envision an even brighter future as we continue to refine and expand this innovative platform.

## Acknowledgments

This study was supported by National Institute of Health (NIH) grants R01AG066653, R01CA266004, R01AG078702, V-Scholar Grant, to R.C.S., R35NS116824 to M.S.G., R35GM142701 to L.C

## Author Contribution Statement

Conceptualization, R.C.S. Methodology, R.C.S., and L.C. and X.M.; Investigation, L H.A.C., C.J.S., T.M., T.R.H., and R.A.R., S.R., J.B., S.B., J.F.A., M.M., B.P., C.W.V., and M.S.G; Writing – Original Draft, R.C.S.; Writing – Review & Editing, R.C.S., H.A.C., C.J.S., S.B., J.F.A., M.M., B.P., C.W.V., M.S.G., and D.B.A.;

Funding Acquisition, R.C.S.; Resources, R.C.S.; Supervision, R.C.S.

## Competing Interest Statement

R.C.S. has research support and received consultancy fees from Maze Therapeutics. R.C.S., M.S.G., and R.C.B. are co-founders of Attrogen LLC. R.C.S. is a member of the Medical Advisory Board for Little Warrior Foundation. M.S.G. has research support and research compounds from Maze Therapeutics, Valerion Therapeutics, Ionis Pharmaceuticals. M.S.G. also received consultancy fee from Maze Therapeutics, PTC Therapeutics, and the Glut1-Deficiency Syndrome Foundation. The remaining authors declare no competing interests

