## Supplementary material for "Spatial Metabolome Lipidome and Glycome from a Single brain Section": sup figure 1-5


**Supplementary Figure and Legends**

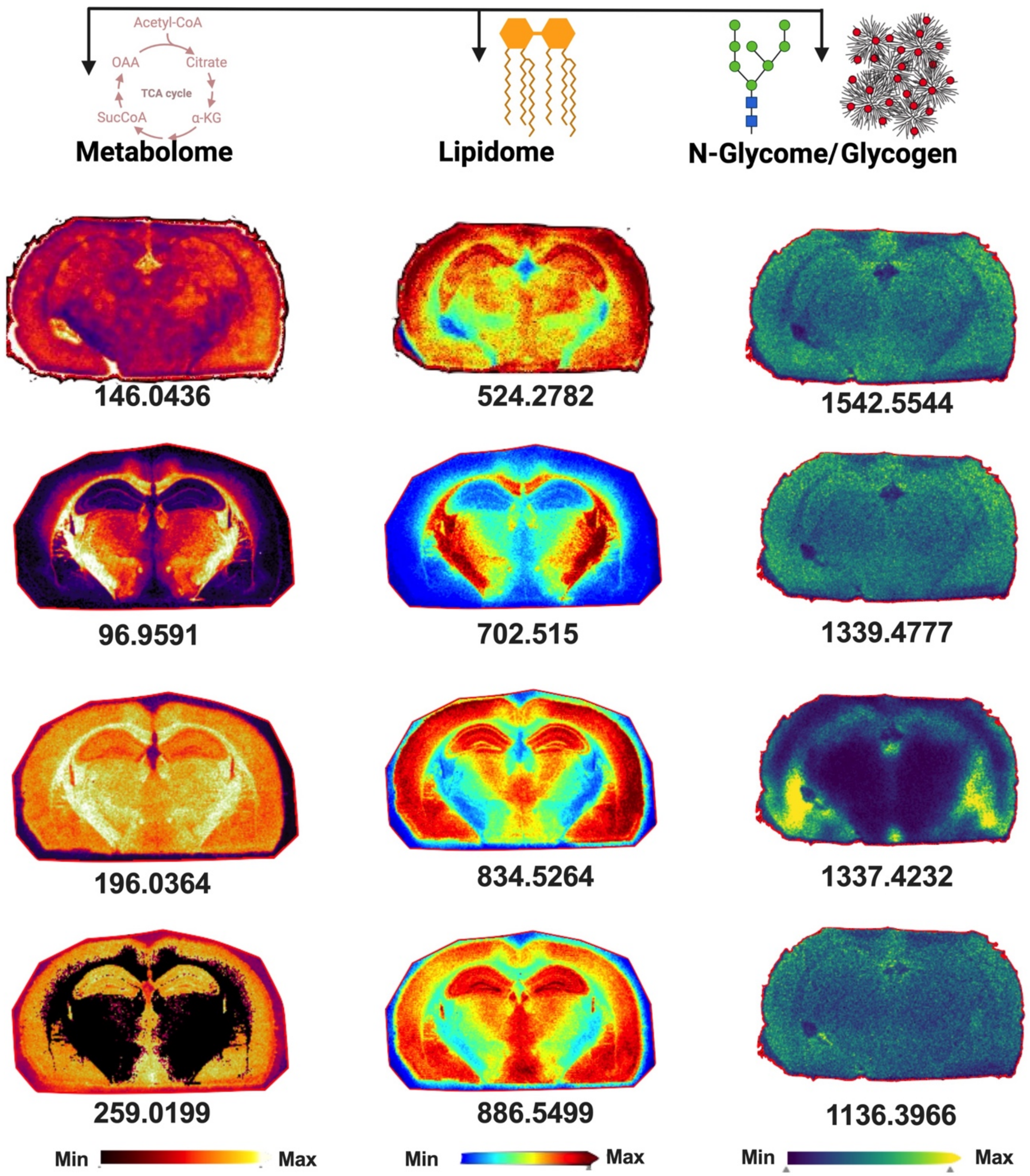


**Supplementary Figure 1. Simultaneous spatial metabolome, lipidome, and glycome by mass spectrometry imaging**

Spatial heatmap images of selected biomolecules from each of metabolome, lipidome, and glycome MALDI imaging runs. All images are produced from the same tissue section following the protocol outlined in figure 1a. *m/z* values are below the heatmap.

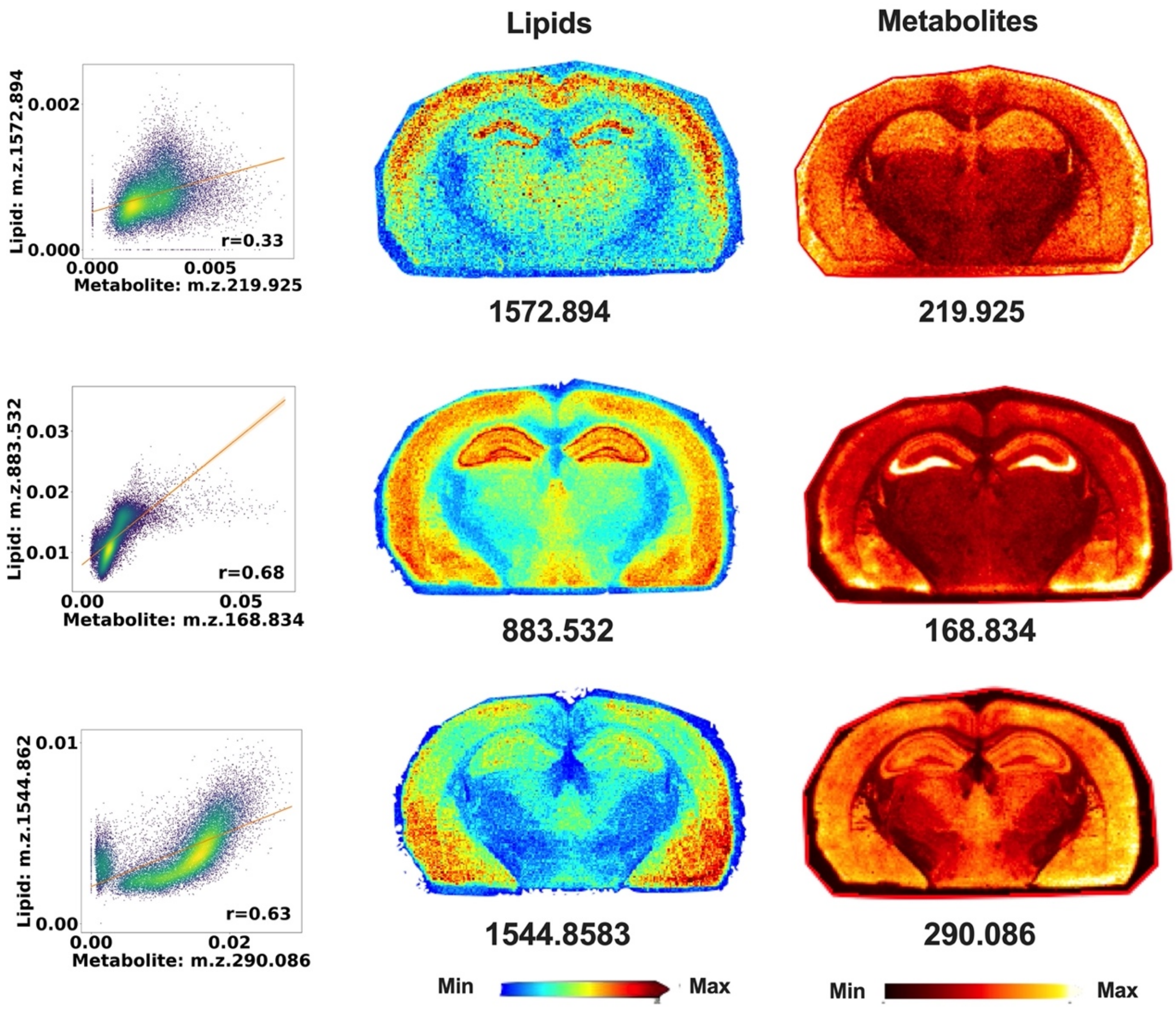


**Supplementary Figure 2. Spatial correlation between lipid and metabolite features after integration.**
Representative paired spatial heatmap images of selected biomolecules from each of metabolome and lipidome MALDI imaging runs based on correlation. representative scatterplots showing correlation between metabolome and lipidome datasets based on Pearson correlation coefficient calculated from the integrated multiome dataset. Representative paired spatial heatmap images of selected biomolecules from each of metabolome and lipidome MALDI imaging runs based on correlation. All images are produced from the same tissue section, *m/z* values are below the heatmap.

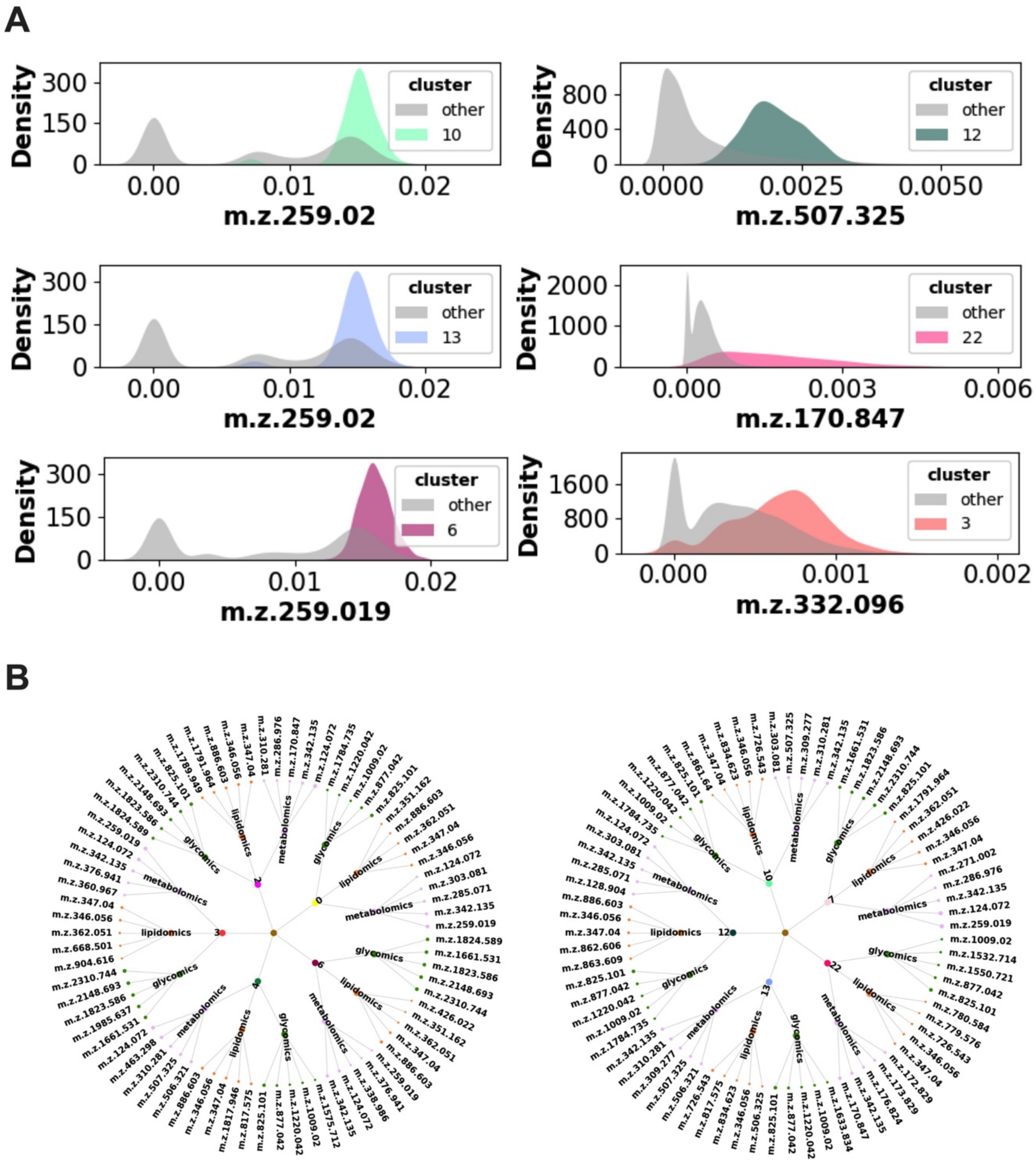


**Supplementary Figure 3. High dimensionality reduction and spatial clustering using different parameters.**

**a** Density plot showing selected overrepresented *m/z* feature in a unique cluster compared to average of rest of the clusters. **b** Chord diagram showing top 5 features from each of the omics analysis among selected clusters.

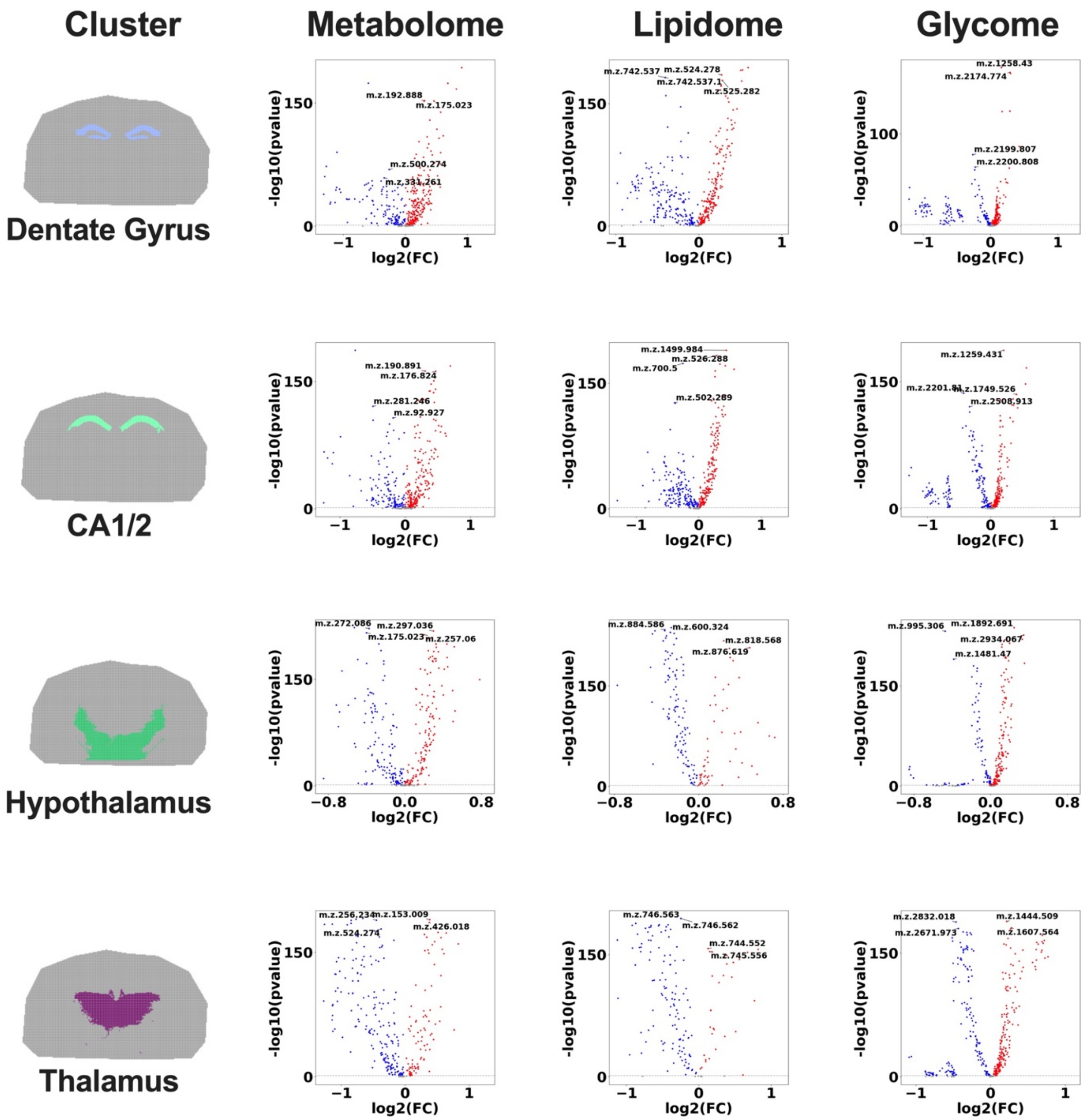


**Supplementary Figure 4. Significantly changed features in unique brain cluster/region.** Spatial map showing only the cluster corresponding to the Dentate gyrus, CA1/2, hypothalamus, and thalamus regions of the brain and their corresponding volcano plot for metabolome, lipidome, and glycome datasets showing significant changes in different brain regions. -$\log_{10}(0.05)$ is the cut for significance. Increased features are in red and decreased features are in blue.

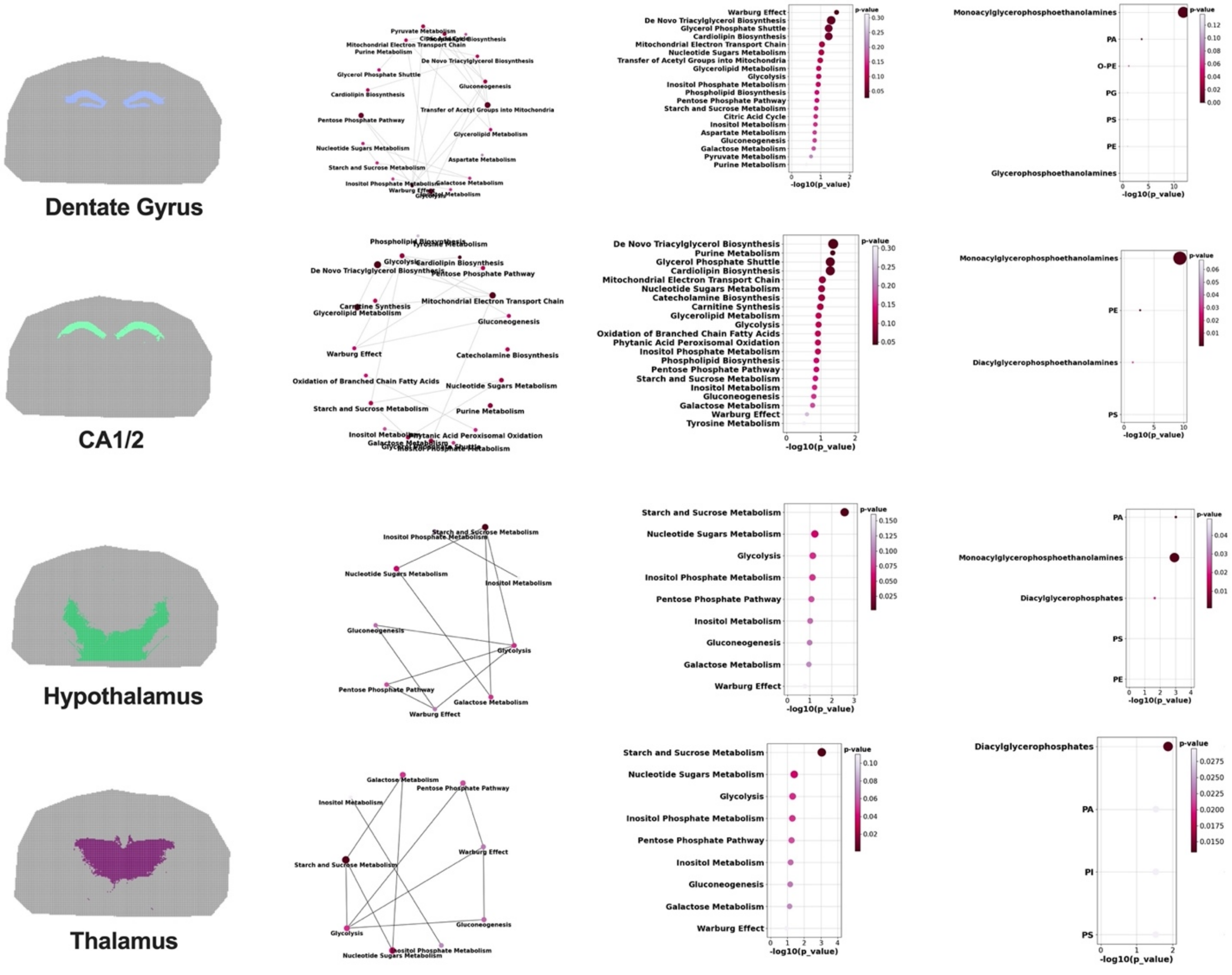


**Supplementary Figure 5. Pathway enrichment within the Sami pipeline.**
Spatial map showing only the cluster corresponding to the Dentate gyrus, CA1/2, hypothalamus, and thalamus regions of the brain and their corresponding metabolic network plot and pathway enrichment based on enriched metabolic pathways generated from the Metaboanlyst 3 R package embeded in Sami.
